# An in silico exploration of combining Interleukin-12 with Oxaliplatin to treat liver-metastatic colorectal cancer

**DOI:** 10.1101/710434

**Authors:** Qing Wang, Zhijun Wang, Yan Wu, David J Klinke

## Abstract

**Background:** Combining anti-cancer therapies with orthogonal modes of action, such as direct cytotoxicity and immunostimulatory, hold promise for expanding clinical benefit to patients with metastatic disease. For instance, a chemotherapy agent Oxaliplatin (OXP) in combination with Interleukin-12 (IL-12) can eliminate pre-existing liver metastatic colorectal cancer and protect from relapse in a murine model. However, the underlying dynamics associated with the targeted biology and the combinatorial space consisting of possible dosage and timing of each therapy present challenges for optimizing treatment regimens. To address some of these challenges, we developed a predictive simulation platform for optimizing dose and timing of the combination therapy involving Mifepristone-induced IL-12 and chemotherapy agent OXP.

**Methods:** A multi-scale mathematical model comprised of impulsive ordinary differential equations was developed to describe the interaction between the immune system and tumor cells in response to the combined IL-12 and OXP therapy. An ensemble of model parameters were calibrated to published experimental data using a genetic algorithm and used represent three different phenotypes: responders, partial-responders, and non-responders.

**Results:** The multi-scale model captures tumor growth patterns of the three phenotypic responses observed in mice in response to the combination therapy against a tumor re-challenge and was used to explore changing the dose and timing of the mixed immune-chemotherapy on tumor growth subjected to a tumor re-challenge in mice. An increased ratio of CD8+ T effectors to regulatory T cells during and after treatment was key to improve tumor control in the responder cohort. Sensitivity analysis indicates that combined OXP and IL-12 therapy worked more efficiently in responders by increased priming of T cells, enhanced CD8+ T cell-mediated killing, and functional inhibition of regulatory T cells. In a virtual cohort that mimics non-responders and partial-responders, simulations show that an increased dose of OXP alone would improve the response. In addition, enhanced IL-12 expression alone or an increased number of treatment cycles of the mixed immune-chemotherapy can barely improve tumor control for non-responders and partial responders.

**Conclusions:** Overall, this study illustrates how mechanistic models can be used for in silico screening of the optimal therapeutic dose and timing in combined cancer treatment strategies.

## Background

Carcinomas of the colon or rectum, termed colorectal cancer, are the third most common cancer diagnosed in both men and women in the United States. The American Cancer Society estimates the number of new cases of colorectal cancer in the United States for 2019 at 145,600 ([1]). With 60,000 fatalities per year, colorectal cancer is second only to lung cancer as a cause of cancer-related deaths in the United States. Upon diagnosis, 10%-20% of patients have already developed liver metastases while 70% of patients with colorectal cancer ultimately develop liver metastases. Unfortunately, the prognosis for patients with liver metastatic colorectal cancer is poor because hepatectomy, palliative chemotherapy and symptomatic treatments are the only available options ([2]).

Interleukin-12 (IL-12) is a potent immunostimulatory cytokine that activates the proliferation and function of key cellular effectors of innate and adaptive immunity such as T lymphocytes and natural killer (NK) cells ([3], [4], [5]). While toxicity is a serious obstacle for use of IL-12 as a systemic therapy in humans, an attractive alternative is to use adenoviral vectors to induce expression in specific tissues. However, transgene expression tends to be transient and the efficacy of re-administration is impaired by the rapid emergence of neutralizing antibodies ([3]). To allow a good control of the strength and duration of IL-12 expression, high-capacity adenoviral vectors containing a liver-specific, Mifepristone-inducible system for the expression of murine IL-12 (HC-Ad/RUmIL-12) were recently designed to control primary or metastatic liver cancer ([6]). Since stand-alone chemo- or radiotherapeutic regimens are insufficient (with a few notable exceptions) to eradicate neoplastic lesion ([7]), HC-Ad/RUmIL-12 was combined with chemotherapy agent Oxaliplatin (OXP) to treat liver-implanted colon cancer cells ([6]). As a consequence of the combination therapy, pre-existing liver metastases of colorectal cancer were eradicated, and enhanced establishment of a protective immune response against tumor rechallenge and increased overall survival of animals were observed. In addition, a dramatic increase in the ratio of cytotoxic CD8^+^ T lymphocytes to immunosuppressive cell populations was detected in the tumor microenvironment ([3]).

Mathematical modeling using systems of ordinary differential equations (ODEs) can improve the design and administration of cancer treatments, especially when experimental data are incorporated ([8], [9], [10], [11], [12], [13]). In silico screening of parameter regions that seem most promising for optimal timing and dosage of therapy can be suggested using calibrated mathematical models and clinical trials can focus on those regions ([13], [14], [15], [16], [17]). For instance, a quantitative systems pharmacology model in [8] was developed to reproduce experimental data of CT26 tumor size dynamics upon administration of RT and an anti-PD-L1 agent in ([18], [19]). The calibrated model was further used as an in silico tool to predict the best treatment combination schedules and sequences. Over the past years, a variety of ODE-based mathematical models have been developed to better understand cancer progression and response to immunotherapy (see details in [20],[21],[22], [23], [24], [25]). In exploring immunotherapy in combination with other treatment modalities, de Pillis et. al developed an ODE model governing cancer growth on a cell population level with a combination of immuno-chemotherapy treatments ([26],[27], [28], [29]). In addition, Kim and colleagues formulated a mathematical model of therapy with oncolytic viruses that simultaneously express immunostimulatory cytokines and costimulatory molecules ([12]). Inspired by these studies, we developed an impulsive ODE model to represent the interaction between tumor and immune system in response to the chemotherapy drug OXP combined with liver-specific expression of IL-12 therapy to explore therapeutic options in the context of liver metastatic colorectal cancer. The current model extends the impulsive ODE model in [13] that only considered an immunotherapy initiated by an adenovirus vaccination to stimulate a tumor-associated antigen-specific T cell response.

The structure of this paper is as follows. First, we present a multi-scale mechanistic model of anti-tumor immunity and tumor growth in response to a combined immuno-chemotherapy using a set of impulsive ODEs. Second, we describe how we calibrated the model parameters against published experimental data using a genetic algorithm. Next we investigate the stability of tumor-free and high tumor equilibria based on the linearized system. Then we study how alter parameter values may change the tumor growth dynamics. Finally, we used the simulation platform to explore potential ways to improve treatment regimes for non-responders and partial responders.

## Methods

Our method was to develop a multi-scale impulsive ODE model based on our understanding of the corresponding biology, which is described in the following paragraphs. Numerical solutions of the model were obtained using simulators generated by C Sharp. The resulting mechanistic mathematical model was calibrated against existing experimental data.

A genetic algorithm was used to find parameter sets that closely match the experimental data in [6]. Each parameter set was modeled using an individual chromosome in order to apply the algorithm to search in the parameter space. For each generation, the impulsive ODE set was solved using the Runge-Kutta method of order four for each individual or parameter set ([30]). The fitness function value, or variance, was calculated using the sum of error squared between experimental data and corresponding model predictions. To reduce the dependence of our model predictions about optimal treatment strategies for the combined therapy on any individual calibrated set of parameter values, we generated an ensemble of 30 parameter sets for each phenotypic cohort (i.e., responder, partial responder, and non-responder) that generated similar good fits against the experimental data. The simulation results using these ensemble of parameter sets were characterized in terms of the mean, 90th percentile upper, and lower predicted responses. Simulations start on day 0, which corresponds to the time of tumor implantation. At the initial time point, we assume that there is no activated tumor specific effector T cells present in the blood and at the site of the tumor. The calibrated mechanistic model was then used to investigate the long-term behavior through stability analysis. Details of model development, parameter calibration, goodness of fit, difference between major variables of immune response of responder mice, partial responder mice, and non-responder mice after the combination treatment, and local stability analysis are discussed in the following subsections.

## Results

### A multi-scale model of tumor growth subject to IL-12 and OXP therapy

Our mathematical model is based on the experimental data presented by Manuela Gonzalez-Aparicio and collaborators in [6] using the MC38Luc1 cell lines for murine metastatic colorectal cancer. Using this mouse model, OXP and IL-12 combination therapy eradicated pre-existing liver metastases, established a protective immune response against tumor rechallenge, and increased overall survival of animals. To better understand the dynamics of the primary response to adenovirus-mediated induction of an anti-tumor immune response, we developed a three-compartment mathematical model to quantify the cytotoxic CD8^+^ T cell response to IL-12 and OXP combined therapy and subsequent inhibition of tumor cell growth, as shown schematically in Figure 1. Among these three compartments, we consider the dynamics of fifteen state variables that are regulated by the following governing biological processes and assumptions:

1. **Naïve CD8^+^ T cells (*T_N_*, units: cells per mm^3^).** We assume that naïve CD8^+^ T cells are produced at a constant rate *c*_1_ from thymus and die naturally at a rate *k*_*d*1_ · *T_N_* ([31]). Naïve T cells are recruited and activated by tumor antigens presented by APC_1_ (antigen-presenting cells in lymph node) at a rate 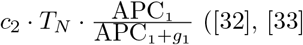 ([32], [33], [34]).

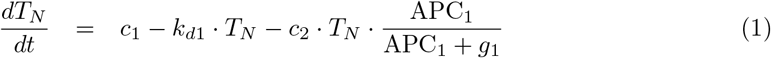
2. **Effector CD8^+^ T cells in lymph node** (*T*_*E*1_, **units: cells per mm**^3^). The increase in the rate of concentration of effector CD8^+^ T cells in the lymph node due to activation of naïve CD8^+^ T cells from the blood stream is given by 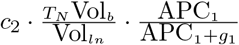, where Vol_*b*_ = 1.4 * 10^3^ *mm*^3^ is the volume of the blood compartment ([35]) and Vol_*ln*_ = 0.25*mm*^3^ is the volume of the lymph node compartment ([36]). We assume that the natural death of effector T cells in the lymph node is negligible. Effector CD8^+^ T cells in the lymph node proliferate at a rate proportional to *T*_*E*1_, a saturable term that represents antigen presenting cells (APC) and defined by 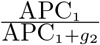, and an immune checkpoint term defined by 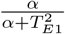, where *α* is the square root of the saturation constant of *T*_*E*1_ ([37]). We also assume that influx rate of effector T cells from blood to lymph node is 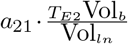, where *T*_*E*2_ is the concentration of T effectors in blood, and *a*_12_ · *T*_*E*1_ is the efflux rate. We assume that *T*_*E*1_ cells are killed by chemotherapy agent OXP_1_ (Oxaliplatin in lymph node) at the rate 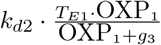.

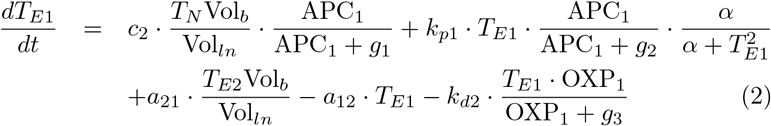
3. **Antigen Presenting Cells in lymph node (APC_1_, units: cells per mm^3^).** We assume that APCs in the lymph node have a natural death rate of *k*_*d*3_ · APC_1_ and the influx rate of APCs from tumor to lymph node is 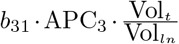, where APC_3_ is the concentration of APCs in tumor, 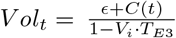 (since *Vol_t_* = *ϵ* + *C*(*t*) + *V_i_* · *T*_*E*3_ · *Vol_t_*, where *T*_*E*3_ is the concentration of T effectors in tumor) is the volume of the tumor compartment, *ϵ* is a small positive constant representing a small volume of tissue that excludes tumor and effector CD8^+^ T cells in the tumor compartment, where *C*(*t*) is the volume of tumor cells in *mm*^3^. The total volume of tumor cells is comprised of the volumes of major histocompatibility complex (MHC) class I positive tumor cells (*C_MHCI^+^_*) and MHC class I negative tumor cells (*C_MHCI^−^_*). The average size of a T effector cell (*V_i_*) is equal to 10^l7^ *mm*^3^ ([38]).

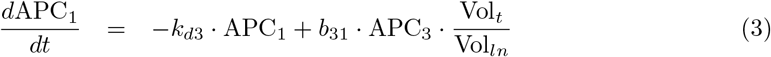
4. **Chemotherapy agent Oxaliplatin in lymph node (OXP_1_, units: mg per kg).** We assume that OXP decays naturally at a rate *k*_*d*4_ · OXP_1_ and the influx rate of Oxaliplatin (OXP) from blood to lymph node is 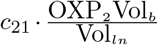, where OXP_2_ is the concentration of OXP in blood.

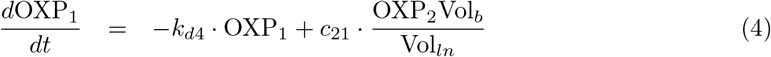
5. **Effector CD8^+^ T cells in blood (*T*_*E*2_, units: cells per mm^3^).** We assume the effector CD8^+^ T cells die naturally at a rate *k*_*d*5_ · *T*_*E*2_ in blood. The influx rate of effector CD8^+^ T cells from lymph node to blood is equal to 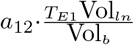 and the efflux rate of effector CD8^+^ T cells from blood to lymph node is equal to *a*_21_ · *T*_*E*2_. The influx rate of CD8^+^ T effectors from the tumor to blood is 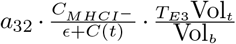, where *T*_*E*3_ is the concentration of T effectors in tumor and the efflux rate of CD8^+^ T effectors from blood to tumor is *a*_23_ · *T*_*E*2_.

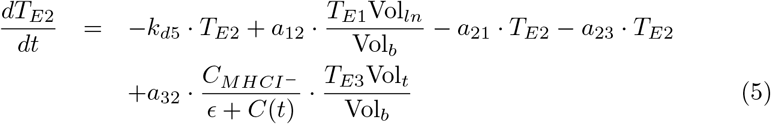
6. **Antigen Presenting Cells in blood (APC_2_, units: cells per mm^3^).** A logistic growth pattern 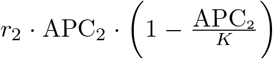 is used for APCs in blood where *r*_2_ is the growth rate constant and *K* is the carrying capacity. We assume a *b*_23_ · APC_2_ efflux rate of APCs from blood to tumor.

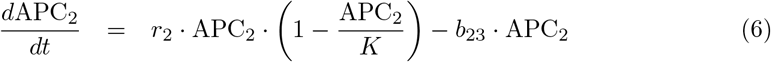
7. **Chemotherapy agent Oxaliplatin in blood (OXP_2_, units: mg per kg).** We assume that OXP decays naturally at a rate *k*_d4_ · OXP_2_ and the efflux rates of OXP from blood to lymph node and from blood to tumor are *c*_21_ · OXP_2_ and *c*_23_ · OXP_2_, respectively. The source of OXP is provided by each administration whose dose and time are reflected by the discrete equation (17).

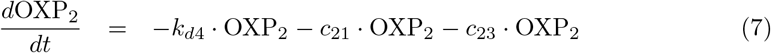
8. **Effector CD8^+^ T cells in tumor microenvironment** (*T*_*E*3_, **units: cells per mm**^3^). We assume that effector CD8^+^ T cells can proliferate locally upon recognition of the corresponding tumor antigen presented by MHC class I positive tumor cells upon IL-12 stimulation and subject to suppression from T regulatory cells at a saturable rate equal to 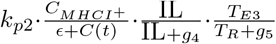, where IL is the concentration of IL-12 and *T_R_* is the concentration of regulatory T cells ([39], [40]). The influx rate of effector CD8^+^ T cells from the blood to tumor is defined by 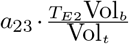. The efflux rate of effector CD8^+^ T cells from the tumor to blood is 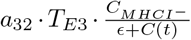. Effector T cells have a finite lifespan and die within the tumor microenvironment at a rate equal to *k*_*d*6_ · *T*_*E*3_. T effector cells in tumor are assumed to be killed by chemotherapy agent OXP in tumor (OXP_3_) at the rate 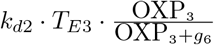.

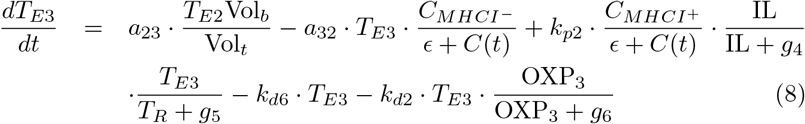
9. **Interferon gamma** (**IFN**_*γ*_, **units: pg per mm**^3^). We assume that IFN_*γ*_ is secreted solely by effector CD8^+^ T cells within the tumor with stimulation from IL-12 and inhibition from regulatory T cells ([41]) at a rate of 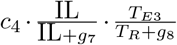. While this assumption may not hold in all model systems, the presence of IFN_*γ*_ in the tumor was dependent on CD8^+^ T cell activation [42]. IFN_*γ*_ decays at a rate proportional to its concentration with a rate constant *k*_*d*7_.

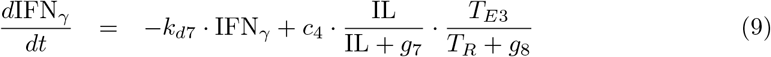
10. **Antigen Presenting Cells in tumor** (**APC**_3_, **units: cells per mm**^3^). We assume that APCs in the tumor microenvironment have a natural death rate of *k*_*d*3_ · APC_3_, the influx rate of APCs from blood to tumor is 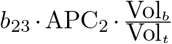 and APCs take tumor antigen in tumor microenvironment and migrate to the lymph node to present tumor antigens to T cells at the rate of *b*_31_ · APC_3_.

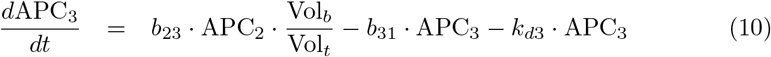
11. **Interleukin-12** (**IL, units: ng per ml**). Interleukin-12 (IL-12) is produced by APCs at a rate of *c*_5_ · APC_3_ and decays naturally at a rate of *k*_*d*8_ ·IL. The extra IL-12 expression obtained through the combined therapy is approximated using a discrete equation (16).

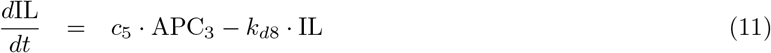
12. **Chemotherapy agent Oxaliplatin in tumor** (**OXP**_3_, **units: mg per kg**). We assume that OXP decays naturally at a rate *k*_*d*4_ · OXP_3_ and the influx rate of OXP from blood to tumor is 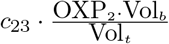.

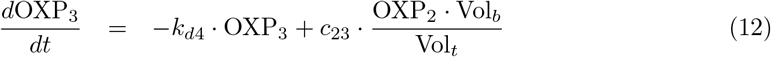
13. **Regulatory T cells** (*T_R_*, **units: cells per mm**^3^). Regulatory T cells are produced at a constant rate *c*_6_ from thymus and die naturally at a rate *k*_*d*9_ ·*T_R_*. We assume that regulatory T cells proliferate at a rate of 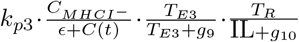 ([3],[4],[6]).

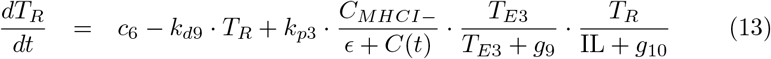
14. **MHC class I positive tumor cells** (*C_MHCI_* +, **units: mm**^3^). MHC class I positive tumor cells are converted from MHC class I negative tumor cells (*C_MHCI_*-) with the assistance of IFN_*γ*_ at a rate 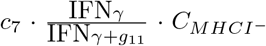 and the rate of effector CD8^+^ T cell-mediated killing of MHC class I positive tumor cells is 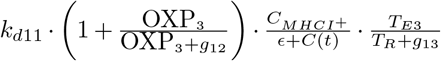 ([6], [31]). We assume that the dilution rate of MHC class I positive tumor cells due to proliferation is *k*_*p*4_ · *C_MHCI_* +. The natural death rate of MHC class I positive tumor cells is assumed to be *k*_*d*10_ · *C_MHCI_* + and MHC class I positive tumor cells are killed by chemotherapy agent OXP in tumor at a rate 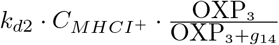.

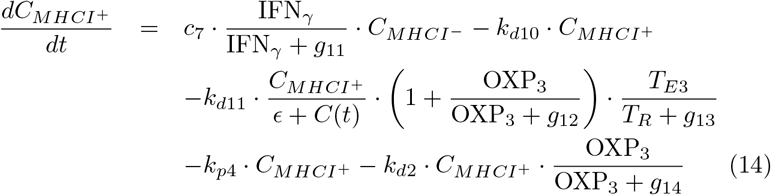
15. **MHC class I negative tumor cells** (*C_MHCI_*-, **units: mm**^3^). MHC class I negative tumor cells are converted to MHC class I positive tumor cells with the assistance of IFN_*γ*_ at a rate of 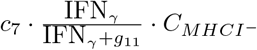. We assume that the proliferation rate of MHC class I positive tumor cells is equal to 2 · *k*_*p*4_ · *C_MHCI_*+. As MHC class I positive tumor cells proliferate, they lose MHC class I expression and become MHC class I negative cells. A logistic growth pattern is assumed for the number of MHC class I negative tumor cells in the absence of treatment. We assume that the natural death rate of MHC class I negative tumor cells is *k*_*d*10_ · *C_MHCI_*- and these cells are killed by chemotherapy agent OXP in tumor at a rate 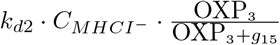. The difference in size of tumor cells caused by tumor rechallenge is described by discrete equation (18).

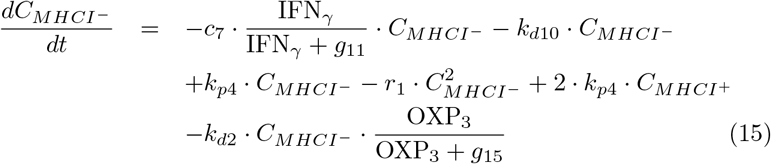
16. We use the following difference equations to reflect the abrupt change of IL-12 concentration and OXP concentration due to therapy and sudden change the size of MHC class I negative tumor due to tumor rechallenge.

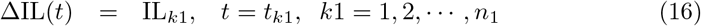

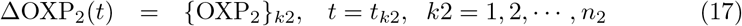

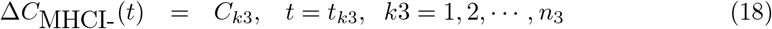

where ΔIL(t) = IL(t^+^) – IL(t^-^) and ΔOXP_2_(*t*) = OXP_2_(*t*^+^) – OXP_2_(*t*^-^) reflect the abrupt changes of IL-12 and oxaliplatin at administration times *t*_*k*1_ and *t*_*k*2_, while IL_*k*_ and {OXP_2_}_*k*2_ are the dosages of IL-12 and oxali-platin at the administration times *t*_*k*1_ and *t*_*k*2_ with *k*1 = 1, 2, …, *n*_1_ and *k*_2_ = 1,2, …, *n*_2_, respectively; Δ*C*_MHCI_-__(*t*) = *C*_MHCI_-__(*t*^+^) – *C*_MHCI_-__(*t*^-^) represents the sudden changes in tumor size due to tumor rechallenge with implantation size *C*_*k*3_ *mm*^3^ at time *t*_*k*3_ with *k*_3_ = 1, 2, …, *n*_3_.

**Figure 1.**
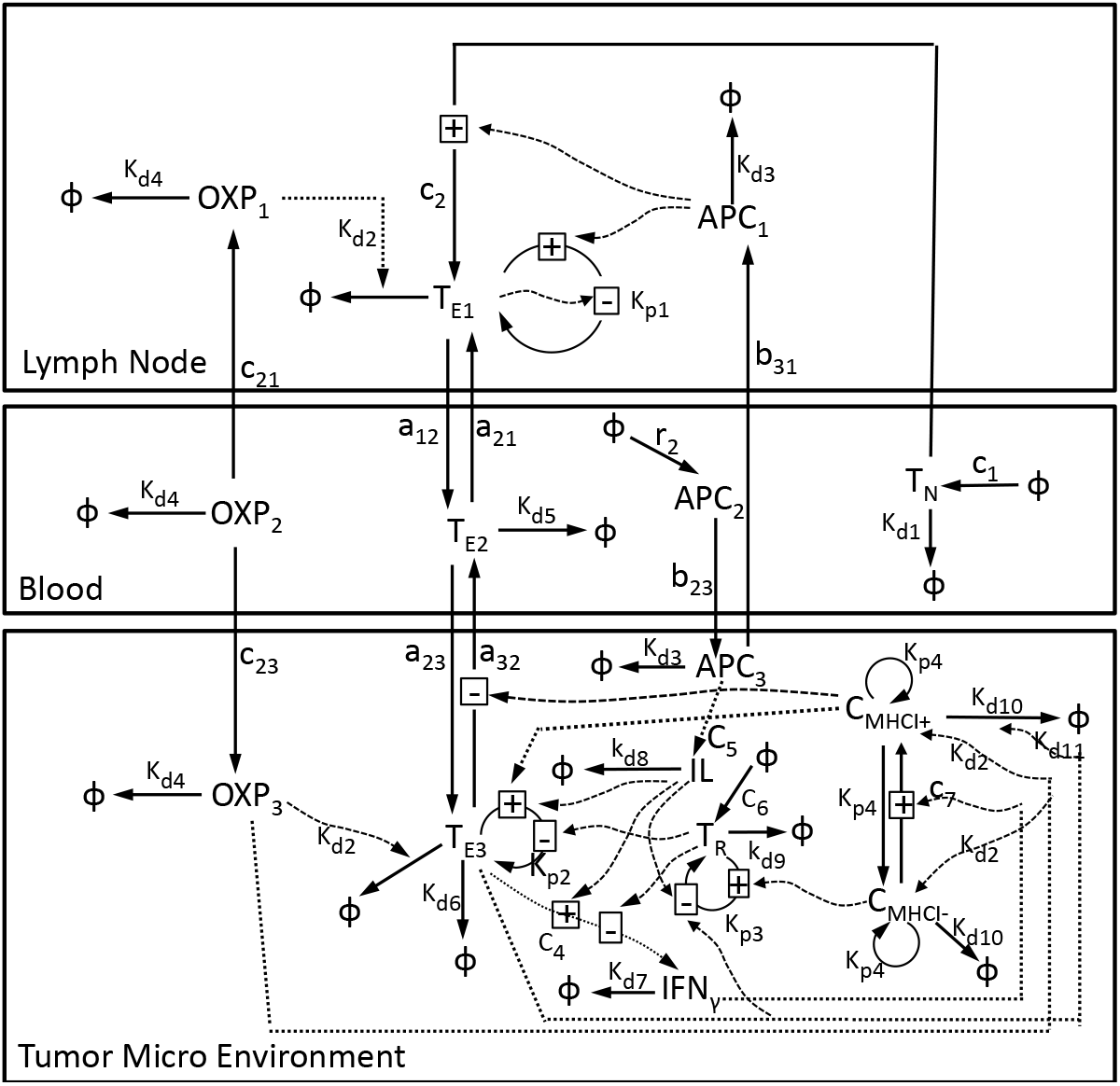
Schematic diagram illustrating the interactions among species present in the three compartments. Naïve CD8^+^ T cells (*T_N_*) are activated and become CD8^+^ T effectors (*T*_*E*1_) when they encounter tumor antigen presented by the antigen presenting cells (APCi) in the lymph node. Once activated, effector CD8^+^ T cells circulate within the blood (*T*_*E*2_) and enter tumor microenvironment (*T*_*E*3_) where they are retained upon recognition of the corresponding tumor-associated antigen. Effector CD8^+^ T cells secrete Interferon gamma (IFN_*γ*_) which assist with the CD8^+^ T cell-mediated killing of tumor cells (*C*_MHCI^+^_ and *C*_MHCI^-^_) through increased presentation of tumor-associated antigens by Major Histocompatibility Complex protein class I (MHCI). During this process, IL-12 (IL) helps promote T cell proliferation and suppresses regulatory T (*T_R_*) cells’ proliferation and immunosuppressing action on effector CD8^+^ T cells. In addition, the chemotherapy drug Oxaliplatin in the lymph node and tumor (OXP_*i*_ where *i* = 1, 3) will kill fast-proliferating cells such as T effectors and tumor cells.

A schematic diagram summarizing this three compartmental model is shown in Figure 1. Model parameters and their meaning are listed in Table 1.

**Table 1.**
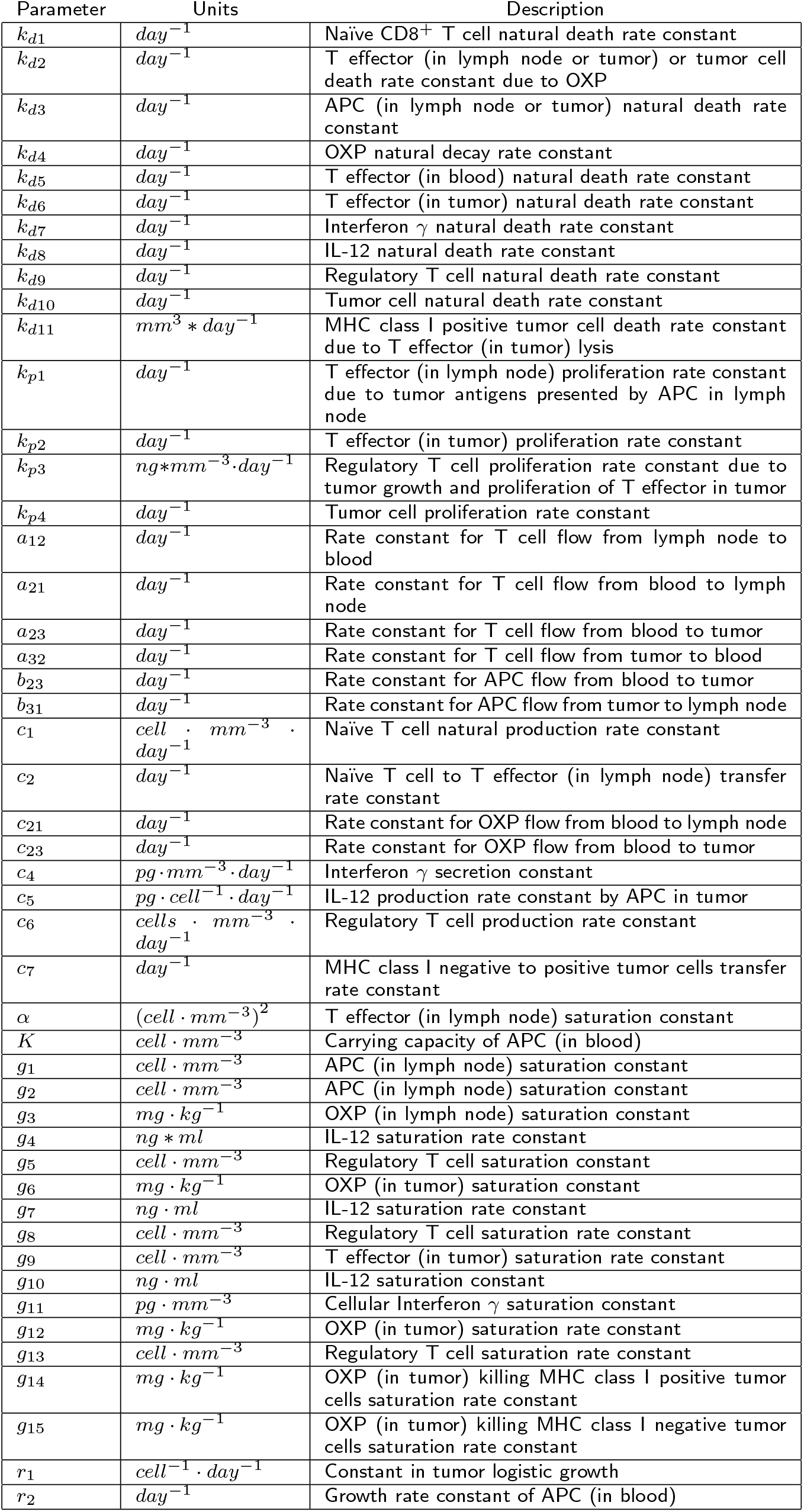
List of parameters in the model

### Non-negativity of solutions to the model

For any mathematical model that has biological implications, it is important to make sure that solutions are non-negative all the time. For our model, we can see that solutions of system comprised of equations 1) – 15) starting from non-negative initial conditions will remain non-negative because 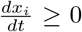 for *x_i_* =0 and *x_j_* ≥ 0 where *i, j* = 1, 2, …, 15 and *i* ≠ *j* with positive impulsive inputs IL_*k*1_, {OXP_2_}_*k*2_, and *C*_*k*3_ at impulsive moments *t*_*k*1_, *t*_*k*2_ and *t*_*k*3_.

### Model calibration

Therapeutic use of IL-12 requires efficient methods to control the plasma concentration of this potent immuno-stimulatory cytokine in order to avoid toxicity ([6]). It was determined in an MC38 syngeneic tumor model that a blood concentration of IL-12 < 20 ng/ml has no anti-tumor effect, while concentrations > 700 ng/ml are associated with toxicity ([43]). Gonzalez-Aparicio and his colleagues designed a new induction protocol to keep IL-12 within this therapeutic range ([6]). Once the liver of a group of C57BL/6 mice was transduced with the vector (typically 2.5*10^8^ IU), a suboptimal amount of Mif (125 *μ*g/kg) is administered for the first 2 days in order to prevent toxicity. The concentration of IL-12 is measured in serum 10 h after the first induction, and based on this information, a stepwise increase in Mif is scheduled to allow several cycles of sustained IL-12 expression in mice treated with the HC-Ad/RUmIL-12 vector (Figure 2). Before we start the calibration of model parameters, we first quantified the IL-12 concentration as a function of time in days (Fig. 2A) and Mifepristone (Fig. 2B) respectively. Empirical functions were used to represent the IL-12 as a function of time and as a function of Mifepristone dose. These calibrated functions are shown in Figure 2, where they are compared against the experimental data reported in ([6]). Overall, the curves show a good match between experimental data and model predictions used to describe Mif-induced IL-12 treatment effects. The authors in [6] verified the Mif-induction system is functional for more than 5 months with a slow decrease in the intensity of expression in each cycle owing to the non-integrative nature of adenoviral vectors. For simplicity, we used the same relationships for each Mif-induced treatment cycle.

**Figure 2.**
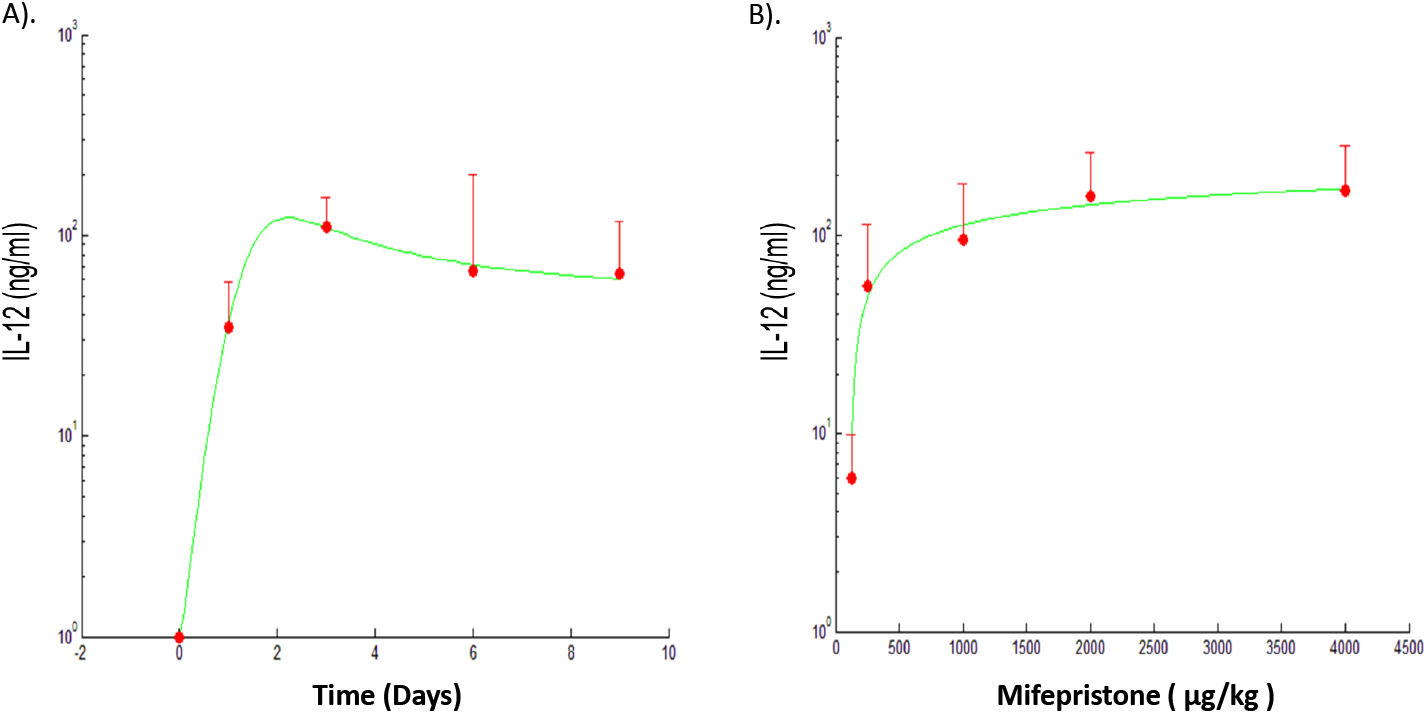
Model fit of Mif-induced IL-12 expression. (A) Simulated IL-12 expression as a function of time was calibrated to (mean + s.d.) experimental data reported in Fig. 1B in ([6]). Specifically, the HC-Ad/RUmIL-12 vector was administered at 2.5*10^8^ IU/mouse in C57BL/6 mice by intrahepatic injection. A set of 8 mice received an adjusted protocol (red circles, n=8) that consisted of 125 *μ*g/kg Mifepristone days 1-2; 250 μg/kg days 3-5; 500 *μ*g/kg days 5-7 and 1000 *μ*g/kg days 9-11. The concentration of IL-12 in serum was determined 10 h after induction at the indicated days. Experimental data in error bars represent mean+ s.d. and the green curve gives our calibrated IL-12 expression (IL) in ng/ml (IL = *f*(*t*) as a function of time in days *t*: 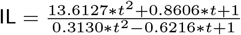. (B) Simulated does-response of IL-12 expression versus Mifepristone dose was calibrated to experimental data in Fig. 1A in [6]. The vector was administered at 2.5*10^8^ IU/mouse in C57BL/6 mice by intrahepatic injection. Two weeks later, a single dose of Mifepristone (125; 250; 1000; 2000 or 4000 *μ*g/kg) was administered intraperitoneally to different groups of animals (n=5). The concentration of IL-12 was measured in serum 10 h later. Experimental data in error bars represent mean+ s.d. and the green curve gives our calibrated IL-12 (IL) expression in ng/ml (IL = *f* (Mif)) as a function of Mifepristone (Mif) in *μ*g/kg Mif: IL = –173.027 + 41.6337 * ln(Mif – 48.0489).

We then calibrated model parameters in system described by equations 1) – 15) using two sets of experimental data from the paper ([6]). The first sets of data are listed below:

1. Total volume of MC38Luc1 tumor cells was calibrated against data shown in Figures 2(B), 2(C), 3 (IL-12 + OXP group).
2. Concentration of Interferon gamma was obtained from Figure 4(B) (IL-12 + OXP group).
3. The ratio between CD8^+^ T lymphocytes and T regulatory cells was obtained from Figure 5(B) (experimental results for IL-12 + OXP group in tumor).

The model was calibrated to these data to reflect the effects of one cycle of Mif-induced IL-12 production combined with chemotherapy drug OXP injection to treat the primary injection of 5 × 10^5^ MC38Luc1 tumor cells into the liver as well as the immunological protection against cancer cells treated with the IL-12 plus OXP combined therapy after a tumor rechallenge with 10^6^ MC38Luc1 tumor cells about 35 days after the completion of the previous combined treatment. Calibration resuits, including the median (solid blue curve), 90th percentile upper (dashed purple curve), and 90th percentile lower responses (dashed green curve) of 30 good fits, are included in Figure 3A), where *C_MHCI^-^_* (0) = 1 *mm*^3^ since *t*_0_ = 0 is the day that 5 × 10^5^ cells/mouse MC38Luc1 tumor cells were inoculated in the liver of C57BL/6 mice ([44, 45]), *T_N_*(0) = 0.0714 cells per *mm*^3^ (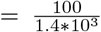 as we assume that the number of naïve CD8^+^ T cells in a mouse is 100 and the volume of the blood system of a mature mouse is 1.4 * 10^3^ *mm*^3^), 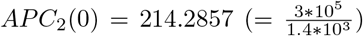 cells per *mm*^3^ according to [46]. Other initial values are zero: *T_Ei_*(0) = *C*_*MHCI*+_ (0) = OXP_*i*_(0) = IFN_*γ*_(0) = APC_1_(0) = APC_3_(0) = T_R_(0) = IL(0) = 0 for *i* = 1, 2,3. Δ*C_MHCI-_* (57) = 2 *mm*^3^, ΔOXP_2_(*s*) = 5 mg/ kg with *s* = 10, 34, 100; ΔIL(t) follows 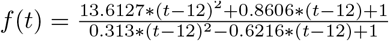 for 12 ≤ *t* ≤ 21 (see details in Fig. 2A). The parameter values used in the simulations are listed in Table 2 with biological meanings of each parameter listed in Table 1.

**Figure 3.**
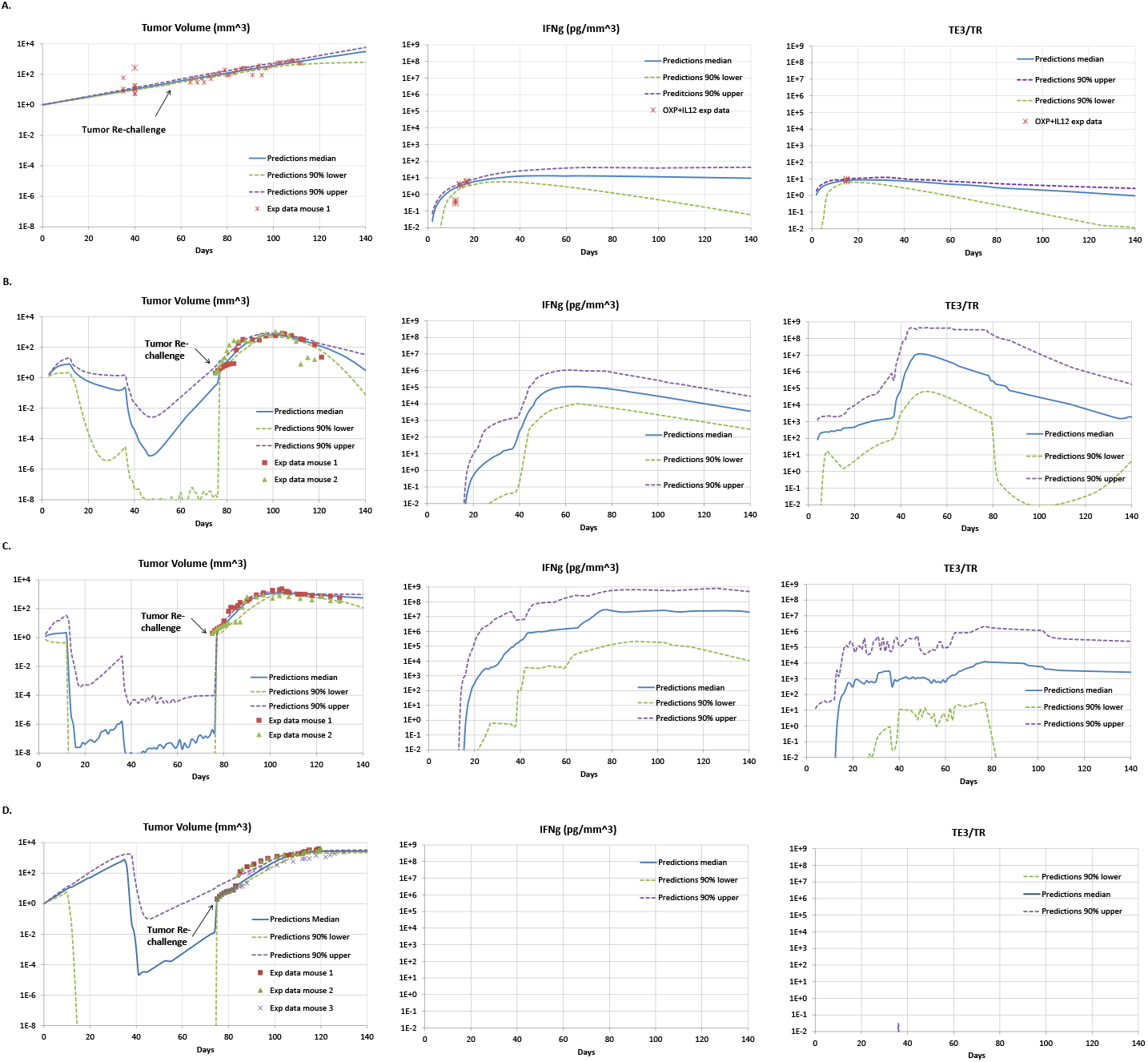
A. Comparison of model predictions with experimental measures of tumor volume, IFNγ and *T*_*E*3_/*T_R_* of mice subjected to tumor rechallenge after one cycle of IL-12 and OXP treatment at day 57. The experimental data were acquired for a group of C57BL/6 mice with 5*10^5^ MC38Luc1 cells inoculated in the liver on day 0 and subjected to one cycle of OXP (on day 9) and Mif-induced IL-12 (started on day 12 and continued 10 days) treatment. To check the immunological protection against cancer cells in treated animals, the cured mice had a tumor rechallenge of 10^6^ MC38Luc1 cells about one month after completion of previous treatment. Experimental measures of tumor volume, IFNγ, and *T*_*E*3_/*T_R_* (crosses) from Figures 2 – 5 in [6] were compared to the model predictions (blue curve) generated using a genetic algorithm. B - D. The experimental data were acquired for a group of C57BL/6 mice bearing hepatic tumors treated with the HC-Ad/RUmIL-12 vector and received two cycles of Mifepristone (Mif) induction preceded by OXP (5 mg/kg, intraperitoneally). Animals cured from their hepatic tumors were subjected to a subcutaneous challenge with the same tumor cells (MC38Luc1), and received a third cycle of IL-12 and OXP treatment starting on day 103. Experimental measures of tumor volume (squares, triangles, and crosses) for mice from Figure 7 in [6] were compared to the model predictions (blue curve) generated using a genetic algorithm. Model predictions calibrated to tumor volume for responder, partial-responder, and non-responder mice treated with one cycle of combined therapy after tumor rechallenge are shown in panels B, C, D, respectively. Each graph displays a collection of 30 good fits of model predictions against experimental data. The solid blue curve provides the median model prediction of the 30 good fits, and the dashed purple and green curves indicate the 90% upper and lower boundaries in the model predictions of 30 good fits, respectively. Example parameter values of good fits in each panel are included in Table 2.

**Table 2.** Examples of calibrated parameter values against experimental data

| Parameter | Fig.3A. | Fig.3B. Par.-Res. | Fig.3C. Res. | Fig.3D. Non-Res. |
| --- | --- | --- | --- | --- |
| $k_{d1}$ | $8.060 * 10^{-2}$ | $4.613 * 10^{-5}$ | $7.892 * 10^{-6}$ | $7.629 * 10^{-5}$ |
| $k_{d2}$ | $3.954 * 10^{-2}$ | 9.923 | 3.286 | 8.370 |
| $k_{d3}$ | $5.738 * 10^{-1}$ | 3.863 | 5.982 | $5.727 * 10^{-1}$ |
| $k_{d4}$ | 1.760 | 2.658 | $1.924 * 10^{-3}$ | 2.358 |
| $k_{d5}$ | $6.293 * 10^{-3}$ | $3.312 * 10^{-4}$ | $5.385 * 10^{-1}$ | $6.170 * 10^{-5}$ |
| $k_{d6}$ | $9.849 * 10^{-3}$ | $5.026 * 10^{-6}$ | $2.258 * 10^{-2}$ | $5.988 * 10$ |
| $k_{d7}$ | $8.159 * 10^{-2}$ | $3.390 * 10^{-2}$ | $7.620 * 10^{-2}$ | 5.603 |
| $k_{d8}$ | $6.097 * 10^{-1}$ | $5.085 * 10^{-6}$ | $4.692 * 10^{-4}$ | $2.513 * 10^{-7}$ |
| $k_{d9}$ | $4.382 * 10^{-6}$ | $5.310 * 10^{-6}$ | $2.206 * 10^{-2}$ | $9.706 * 10^{-7}$ |
| $k_{d10}$ | $2.116 * 10^{-5}$ | $2.140 * 10^{-6}$ | $7.795 * 10^{-6}$ | $9.434 * 10^{-6}$ |
| $k_{d11}$ | $7.956 * 10^{-3}$ | $6.953 * 10^{-6}$ | $7.047 * 10^{-5}$ | $6.539 * 10^{-5}$ |
| $k_{p1}$ | 2.188 * 10 | $4.770 * 10^4$ | 9.808 * 10 | $5.515 * 10^2$ |
| $k_{p2}$ | $6.445 * 10^{-6}$ | $8.367 * 10^{-7}$ | $1.687 * 10^{-12}$ | $9.255 * 10^{-7}$ |
| $k_{p3}$ | $5.015 * 10^{-7}$ | $7.807 * 10^{-9}$ | $4.276 * 10^{-7}$ | $8.696 * 10^{-7}$ |
| $k_{p4}$ | $5.800 * 10^{-2}$ | $3.952 * 10^{-1}$ | $3.297 * 10^{-1}$ | $2.186 * 10^{-1}$ |
| $a_{12}$ | $5.497 * 10^{-1}$ | 9.636 | $3.031 * 10^{-2}$ | $8.575 * 10^{-2}$ |
| $a_{21}$ | $1.133 * 10^{-4}$ | $9.984 * 10^{-1}$ | $9.011 * 10^{-4}$ | $1.614 * 10^{-1}$ |
| $a_{23}$ | $7.254 * 10^{-7}$ | $9.567 * 10^{-6}$ | $2.431 * 10^{-3}$ | $4.776 * 10^{-11}$ |
| $a_{32}$ | $4.575 * 10^{-3}$ | $9.246 * 10^{-1}$ | $2.573 * 10^{-3}$ | $7.638 * 10^{-3}$ |
| $b_{23}$ | $9.581 * 10^5$ | $6.334 * 10^{-9}$ | $1.406 * 10^{-2}$ | $5.710 * 10^{-13}$ |
| $b_{31}$ | $6.872 * 10^{-7}$ | $8.489 * 10^{-8}$ | $5.700 * 10^{-10}$ | $3.980 * 10^{-9}$ |
| $c_1$ | $8.563 * 10^{-4}$ | 9.611 | $4.366 * 10^{-4}$ | $4.108 * 10^{-2}$ |
| $c_2$ | $7.360 * 10^{-2}$ | $8.751 * 10^{-2}$ | $3.373 * 10$ | $6.011 * 10^{-2}$ |
| $c_{21}$ | $7.211 * 10^{-8}$ | $5.684 * 10^{-4}$ | $5.475 * 10^{-2}$ | $9.907 * 10^{-1}$ |
| $c_{23}$ | $3.648 * 10^{-6}$ | 5.873 | 3.355 | $9.905 * 10^{-1}$ |
| $c_4$ | $6.878 * 10^{-2}$ | $8.671 * 10^4$ | $7.547 * 10^2$ | $2.327 * 10^2$ |
| $c_5$ | $5.263 * 10^5$ | $9.844 * 10^{-9}$ | $7.279 * 10^{-10}$ | $9.722 * 10^{-11}$ |
| $c_6$ | $1.490 * 10^{-2}$ | $3.260 * 10^{-2}$ | $6.253 * 10^{-4}$ | $7.635 * 10^2$ |
| $c_7$ | $8.842 * 10^2$ | 7.546 | $3.412 * 10^2$ | $9.289 * 10^3$ |
| $\alpha$ | $5.530 * 10^9$ | $3.647 * 10^6$ | $1.556 * 10^7$ | $9.834 * 10^3$ |
| $K$ | $7.539 * 10^4$ | $9.292 * 10^{12}$ | $3.116 * 10^{11}$ | $4.964 * 10^5$ |
| $g_1$ | $7.971 * 10^9$ | 4.578 | $9.304 * 10^2$ | $3.445 * 10^4$ |
| $g_2$ | $2.441 * 10^{-2}$ | $3.568 * 10^{-10}$ | $6.108 * 10^{-13}$ | $5.108 * 10^{-12}$ |
| $g_3$ | $3.561 * 10^{-7}$ | $1.081 * 10^{-11}$ | $9.294 * 10^{-12}$ | $5.325 * 10^{-5}$ |
| $g_4$ | $6.740 * 10^2$ | $9.535 * 10^4$ | $1.518 * 10$ | $3.713 * 10$ |
| $g_5$ | $9.702 * 10^7$ | $2.440 * 10^5$ | $9.131 * 10^{-3}$ | $8.235 * 10^{11}$ |
| $g_6$ | $2.318 * 10^5$ | $8.034 * 10^3$ | $3.843 * 10^6$ | $6.932 * 10^4$ |
| $g_7$ | $9.387 * 10^7$ | $9.307 * 10^8$ | $9.040 * 10^8$ | $7.528 * 10^8$ |
| $g_8$ | $8.412 * 10^{-5}$ | $7.079 * 10^{-9}$ | $7.897 * 10^{-7}$ | $5.385 * 10^{-8}$ |
| $g_9$ | $3.757 * 10^{-5}$ | $5.234 * 10^{-3}$ | $5.185 * 10^{-8}$ | $9.378 * 10$ |
| $g_{10}$ | $6.242 * 10^{-5}$ | $9.338 * 10^{-6}$ | $5.537 * 10^{-6}$ | $5.110 * 10^{-8}$ |
| $g_{11}$ | $2.183 * 10^8$ | $5.064 * 10^{10}$ | $7.624 * 10^6$ | $4.565 * 10^9$ |
| $g_{12}$ | $3.229 * 10^{-8}$ | $1.542 * 10^{-11}$ | $1.930 * 10^{-10}$ | $7.116 * 10^{-9}$ |
| $g_{13}$ | $8.733 * 10^6$ | $2.273 * 10^7$ | $4.689 * 10^7$ | $9.747 * 10^5$ |
| $g_{14}$ | $5.981 * 10^3$ | $4.809 * 10^9$ | $4.799 * 10^{10}$ | $3.855 * 10^4$ |
| $g_{15}$ | $3.068 * 10^9$ | $7.927 * 10^5$ | $8.642 * 10^3$ | $5.392 * 10^6$ |
| $r_1$ | $2.140 * 10^{-10}$ | $2.448 * 10^{-2}$ | $6.718 * 10^{-2}$ | $7.260 * 10^{-5}$ |
| $r_2$ | 8.065 | $2.780 * 10^{-4}$ | $3.394 * 10^{-9}$ | $2.793 * 10^{-6}$ |

To show the long-term management of colorectal cancer using the combined therapy, tumor growth of a group of mice subjected to one cycle of treatment was calibrated to data from Figure 7(D) in [6]. In the combined therapy, 5 mg/kg OXP on day 100 and 10-day IL-12 induction starting day 103, which follows the adjusted protocol as described in Figure 2, were administered after a tumor rechallenge on day 75. This treatment occurred about two weeks after the mice survived two cycles of combined treatments with 10-day Mif-induced IL-12 (induction started on days 13 and 37) and OXP treatments (5mg/kg on days 10 and 34), which in turn started about two weeks after the initial implantation of MC38Luc1 tumor cells on day 0. The experimental results were split into responders (Fig. 3B), partial responders (Fig. 3C) and non-responders (Fig. 3D) groups. The mathematical model was calibrated separately to these different response groups. Figures 3B – 3D with the median (solid blue curve), 90th percentile upper (dashed purple curve), and 90th percentile lower responses (dashed green curve) of 30 good fits illustrate the results of our simulations compared with the corresponding experimental data. Here, we have Δ*C_MHCI-_* (75) = 2 *mm*^3^, ΔOXP_2_(9) = 5 mg/kg; Δ*IL*(*t*) follows 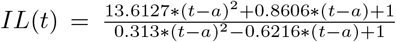 for *a* ≤ *t* ≤ *a* + 9 with *a* = 13, 37, 103. A sample set of parameter values for each of the response groups of mice used in the simulations are listed in Table 2.

*Difference in treatment efficacy: non-responders versus responders* While both the responders and non-responders survived two cycles of combination therapy treatment before tumor rechallenge and then underwent the third cycle following the tumor rechallenge on day 75, the simulations in Figs. 3B – 3D suggest that the nonresponders show near zero concentration of IFN_*γ*_ and near zero ratio of T effectors to regulatory T cells in the tumor all the time comparing to a stable concentration of IFN_*γ*_ and ratio of T effectors to regulatory T cells in the tumor (at least 10^3^ cells per *mm*^3^ after the combination therapy treatment) in responders and partial responders. The simulations also indicate that whether the immune system can maintain a high ratio of T effectors to regulatory T cells in the tumor as well as generating a moderate but stable concentration of IFN_*γ*_ might be crucial to control tumor growth. This finding is consistent with results from experimental studies ([7], [47]).

### Goodness of fit

Descriptions of model parameters are shown in Table 1. A couple of sample sets of estimated values of parameters obtained through fitting predictions of the impulsive ODE model 1) – 18) to data from a group of experiments in [6] are listed in Table 2. Figure 3A illustrates the comparison between model solutions and experimental measurements on tumor control and immunological protection against cancer cells in animals treated with IL-12 plus OXP. Experimental results for long-term management of colorectal cancer by observing cooperation of IL-12 and OXP for the control of experimental relapses in distant locations are compared against model predictions in Figures 3B, 3C, and 3D for responders, partial-responders and nonresponders, respectively. Trajectories of tumor growth, IFN_*γ*_ and ratio of *T*_*E*3_ to *T_R_* arising from the model are extremely close to corresponding data from the experiments. For each calibration, an excess of data points (57, 65, 58, 84 for Figures 3A – 3D, respectively) relative to the number of parameters (48) suggests that the model is identifiable in theory.

## Model stability analysis

In this section, we discuss local stability of equilibria of the model using linearized system evaluated at these points. We found that system comprised of equations (1) – (15) has a tumor-free equilibrium 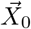, a second tumor free equilibrium 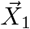 when APC_2_ growth rate is larger than the rate constant for APC2 flowing from blood to tumor (i.e., r_2_ > b_23_), and a high tumor equilibrium 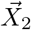 when proliferation rate of tumor cells is higher than natural death rate of tumor cells (i.e., *k*_*p*4_ > *k*_*d*10_).

By setting the right hand sides of the equation system (1) – (15) to zero and solving the equations simultaneously, we obtain

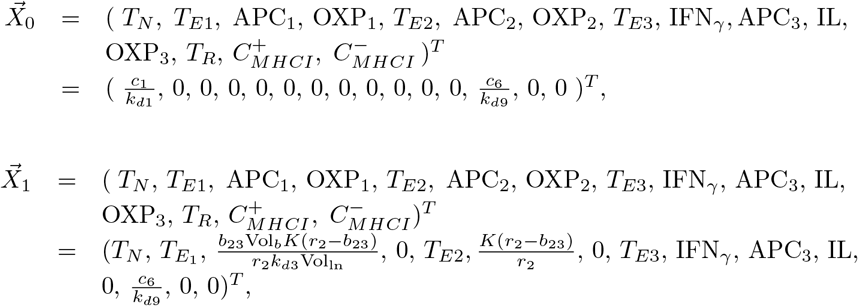

where *T*_*E*1_ satisfies the following polynomial equation

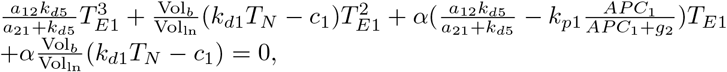

(Based on the Descartes’ rule, there is only one positive solution from the equation), 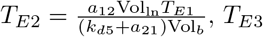 satisfies the following quadratic equation

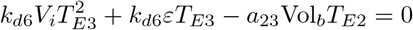

which has only one positive solution 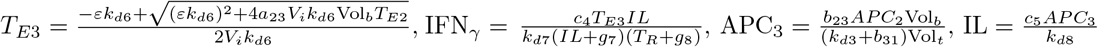 and

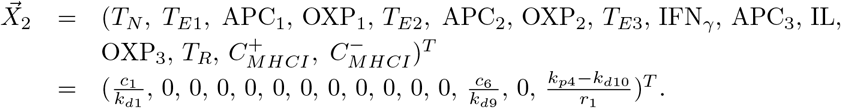

By simple calculation, thirteen of the fifteen eigenvalues of the Jacobian matrix of linearized system at the equilibrium 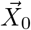 are listed below

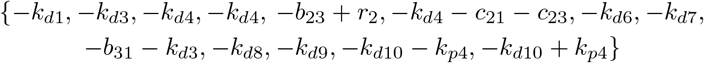

The rest of the eigenvalues are obtained from solving the following quadratic equation:

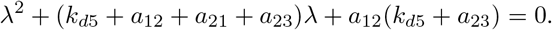

It is easy to see that both eigenvalues are in the left half of the complex plane for a wide range of the parameters. Thus the first tumor free equilibrium 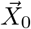 is stable if both *r*_2_ < *b*_23_ (i.e., APC_2_ growth rate is smaller than the rate constant for APC_2_ flowing from blood to tumor) and *k*_*p*4_ < *k*_*d*10_ (i.e., tumor proliferation rate is less than tumor natural death rate), otherwise it is unstable.

The second tumor free equilibrium 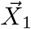 exists when *r*_2_ > *b*_23_ (i.e., APC_2_ growth rate is larger than the rate constant for APC_2_ flowing from blood to tumor). Similar to the previous case, most of the eigenvalues of the Jacobian matrix at 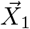 are in the left-half of the complex plane. It is easy to see that twelve of the fifteen eigenvalues are negative. With respect to the remaining three, one is *b*_23_ — *r*_2_, and the other two satisfies the following quadratic equation

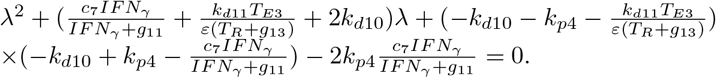

It is found that both eigenvalues are in the left-half of the complex plane if *k*_*p*4_ < *k*_*d*10_. Hence this tumor-free equilibrium point is locally stable if *r*_2_ > *b*_23_ and *k*_*p*4_ < *k*_*d*10_; otherwise it is unstable.

The high tumor equilibrium 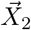 exists when *k*_*p*4_ > *k*_*d*10_. Twelve of the fifteen eigenvalues of the Jacobian matrix are found as

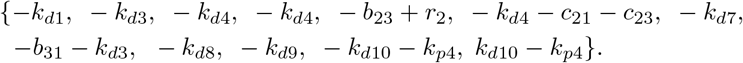

The other three eigenvalues are the roots of the polynomial *p*(λ) = λ^3^ + *α*_1_λ^2^ + *α*_2_λ + *α*_3_, where

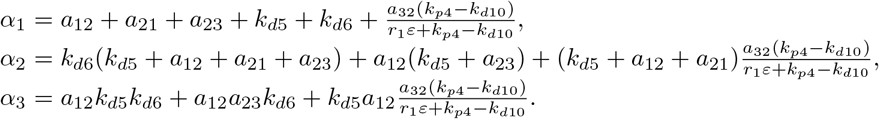

Based on the list of the first twelve eigenvalues, the high-tumor equilibrium is unstable if either *r*_2_ > *b*_23_ or *k*_*d*10_ > *k*_*p*4_ is satisfied. Suppose *r*_2_ < *b*_23_ and *k*_*d*10_ < *k*_*p*4_, it is easy to see that *α*_1_ > 0, *i* = 1, 2, 3. According to the Routh-Hurwitz criterion, *p*(λ) is a Hurwitz polynomial if and only if *α*_1_ *α*_2_ — *α*_3_ > 0, which is indeed the case after simplifying the expression. Therefore, the high-tumor equilibrium is stable if *r*_2_ < *b*_23_ (i.e., APC_2_ growth rate is smaller than the rate constant for APC_2_ flowing from blood to tumor) and *k*_*d*10_ < *k*_*p*4_ (i.e., tumor proliferation rate is larger than tumor natural death rate), otherwise it is unstable.

The stability conditions for tumor free equilibrium 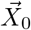 indicate that small tumors may not grow into a threatening size when tumor proliferation rate is less than tumor natural death rate (*k*_*p*4_ < *k*_*d*10_) without any treatment (*r*_2_ < *b*_23_ with all the APC_*i*_, OXP_*i*_, IL and *T_Ei_, i* = 1, 2, 3 in 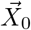 be to zero). Under the combined IL-12 and OXP treatment, MHC class I positive tumor cells will ultimately be eliminated. Depending on effects of the combined treatment (reflected by the remaining level of APC_*i*_ and *T_Ei_, i* = 1, 2, 3 and whether *r*_2_ > *b*_23_), MHCI negative tumor cells (*C_MHCI-_*) will either be completely removed in which case solutions of this dynamic approach the second tumor-free equilibrium 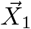 or the MHC class I negative tumor cells eventually approach the carrying capacity. This can occur when MHC class I positive tumor cells are all killed by tumor infiltrating lymphocytes, which results in the exhaustion of effector CD8^+^ T cells and cytokines while naïve T cells and MHC class I negative tumor cells remaining at constant levels.

## Sensitivity of parameters

To test the impact of how the change of a certain parameter value (e.g. *α*) would affect tumor growth pattern for responders, partial-responders, and non-responders, normalized differences of tumor sizes 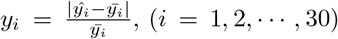 on day 120 (8 days post the second treatment cycle of the combination therapy after tumor rechallenge) were used to draw the violin plots ([48]) in Figure 4 for each of the 48 parameters for all three patient groups, where 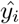 is the predicted tumor size in *mm*^3^ using 0.1 × *α_i_* and other parameters in the ith calibrated parameter set and *y_i_* is the predicted tumor size in *mm*^3^ using *α_i_* and other parameters in the *i_th_* calibrated parameter set and *α_i_* is the calibrated value for parameter *α* in the *i_th_* calibrated parameter set.

**Figure 4.**
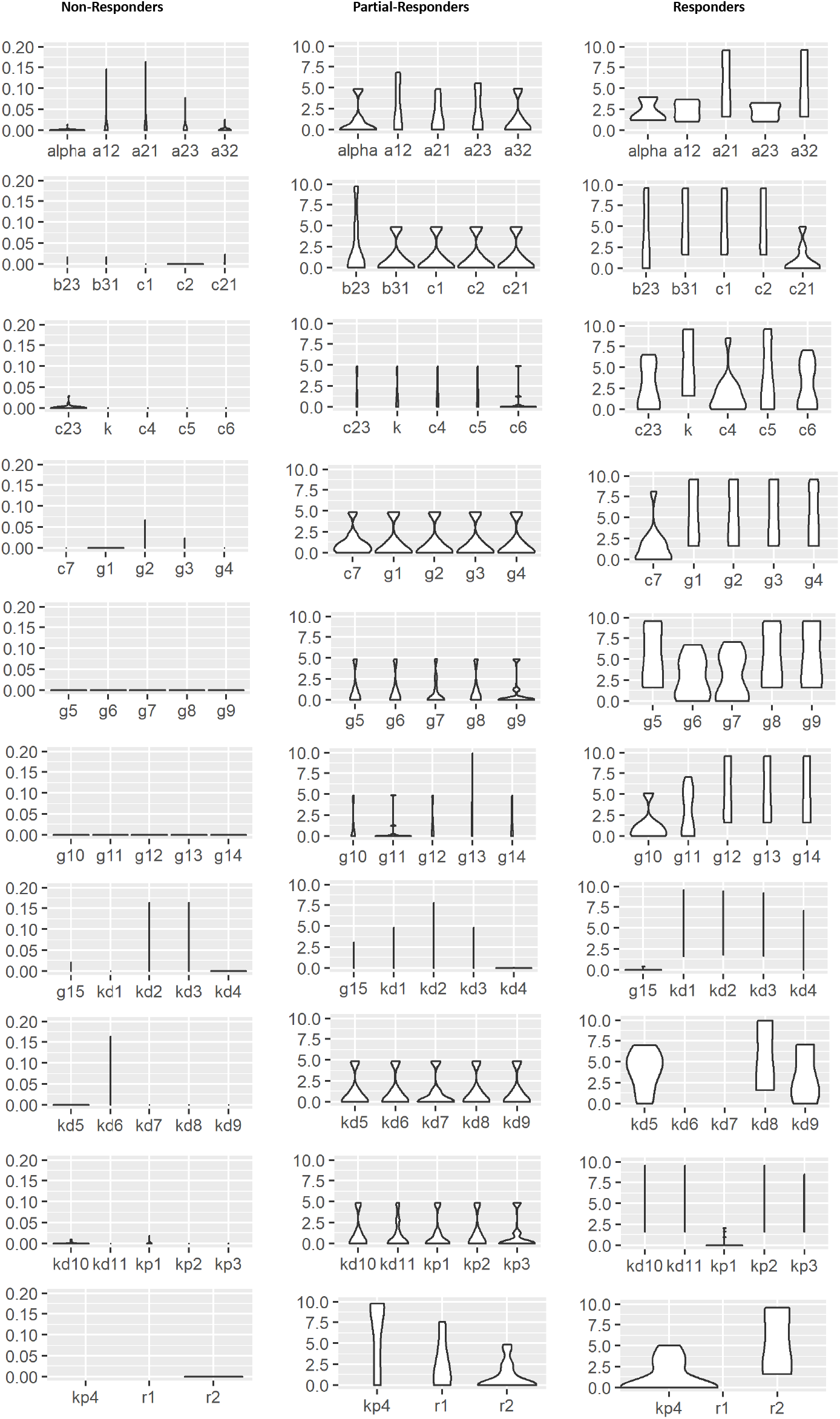
Violin plots of normalized tumor size changes with 30 good fits of parameter sets for responders, partial-responders, and non-responders on day 120. The sample set of parameter values for each group used in the plots are listed in Table 2.

In general, we found that changing the value of each of the 48 parameters barely affected the tumor growth for non-responders. In addition, there are 10 (out of 48) parameters whose value changes greatly affect tumor size of responders but not the size of non-responders and partial-responders. These potentially OXP and IL-12 treatment important parameters include *c*_23_ (OXP flow rate from blood to tumor), *K* (APC carrying capacity), *c*_4_ – *c*_6_ (IFN_*γ*_, IL-12, and T_*R*_ production rate constants, respectively), *g*_10_ – *g*_14_ (IL-12, IFN_*γ*_, OXP_3_, T_*R*_, and 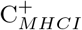 killing by OXP_3_ saturation rate constants, respectively). In addition, changes in the value of the following 7 parameters cause from zero for non-responders to increasing changes in normalized tumor size from partial-responders to responders: T cell flow rates from blood to lymph node and from tumor to blood, *a*_21_ and *a*_32_, respectively; APC flow rates from tumor to lymph node *b*_31_; production rate constant of naive T cells *c*_1_; transfer rate constant of naive T cell to T effector cell in lymph node *c*_2_; IL-12 natural death rate constant *k*_*d*8_, and APC growth rate constant *r*_2_. We also note that no change in tumor size for all three mice group (non-responders, partial-responders, and responders) results from the value changes of following 5 parameters: 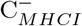 killing by OXP_3_ saturation rate constant g_15_, natural death rate constant of naive T cells *k*_*d*1_, killing rate constant of T effectors or tumor cells by OXP *k*_*d*2_, APCs natural death rate constant *k*_*d*3_, and natural decay rate constant of OXP *k*_*d*4_ (see Fig. 4).

## Model simulations

In order to investigate potential ways to improve treatment regimes for partial-responders and non-responders, we simulated the following alternative treatment scenarios: changing the dose and frequency of chemotherapy drug OXP administration, changing the strength of Mif-induced IL-12 expression, and changing the number of combined treatment cycles.

### Partial-responders

We note from Figure 5 that increased number of treatment cycles in the IL-12 and OXP combination therapy does not seem to improve tumor control in the first 8 months post treatment while increased dose of OXP alone would achieve better tumor control and enhanced strength of IL-12 expression alone would slightly reduce tumor size more rapidly after the tumor reaches its maximum size.

**Figure 5.**
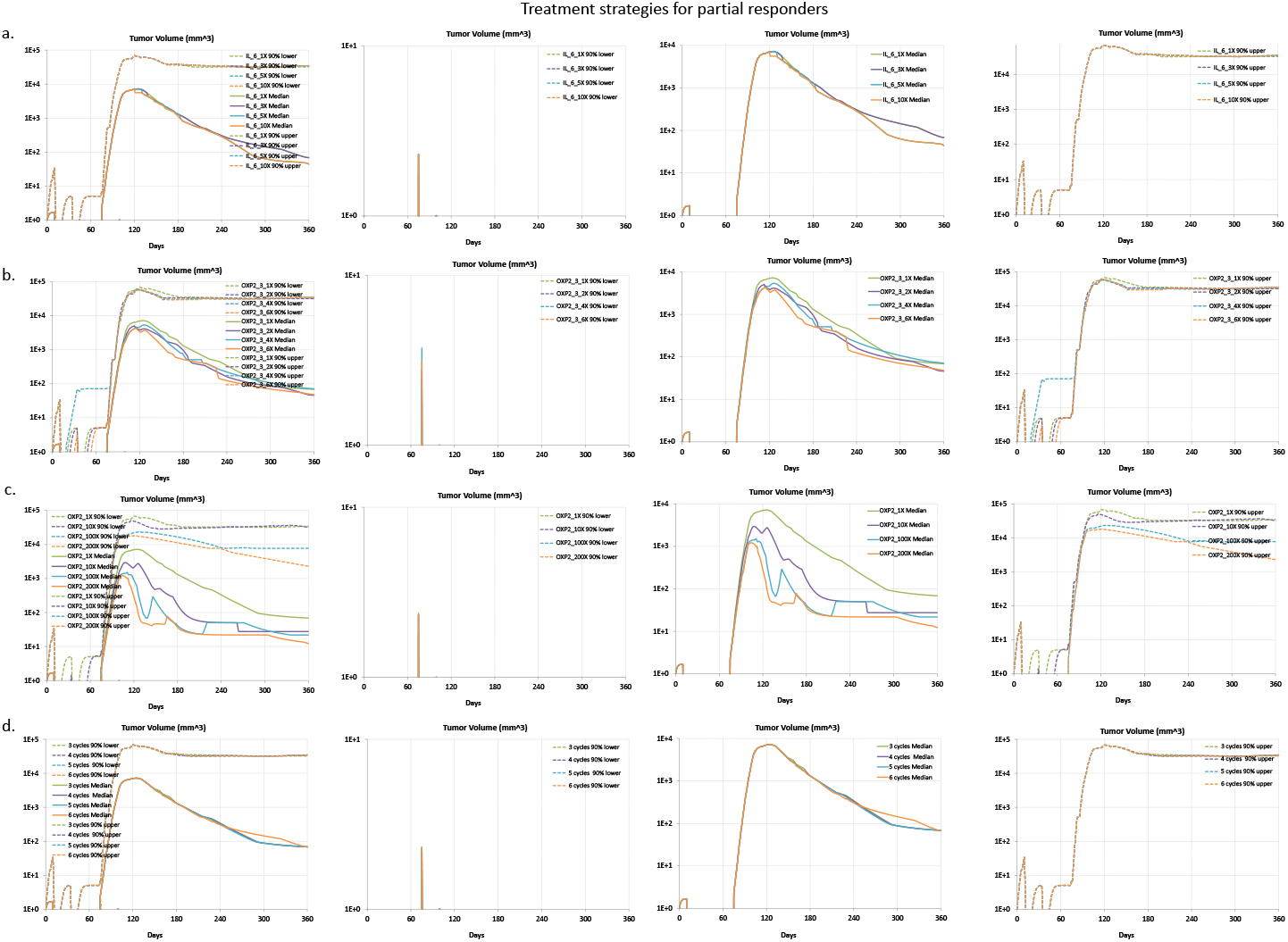
Treatment strategies for partial-responders. The median, 90th percentile lower, and 90th percentile upper responses of 30 patients were sketched for each treatment strategy using 30 sets of good fits of calibrated parameters for partial responders. See a sample set of parameter values used in the plots in Table 2. a. Effects of increased interleukin-12 (IL-12) dose from 1 time (1X, control) to 3, 5, 10 times (3X, 5X, 10X, respectively). b. Effects of moderately increased OXP dose from 1 time (1X, control) to 2, 4, 6 times (2X, 4X, 6X, respectively). c. Effects of aggressively increased OXP dose from 1 time (1X, control) to 10, 100, 200 times (10X, 100X, 200X, respectively). d. Effects of increased number of treatment cycles from 3 cycles to 4, 5, and 6 cycles.

### Non-responders

From Figure 6, we found that neither increased strength of IL-12 expression nor moderately increased OXP dose alone in the IL-12 and oxaliplatin (OXP) combination therapy seems to improve tumor control for the median, 90th percentile lower and 90th percentile upper responses for the 30 non-responder patients. Meanwhile, aggressively increased OXP dose (for instance, 10+ times) in the combination therapy shows reduced tumor size and delayed time of tumor reaching its carrying capacity only for the 90th percentile lower responses for the 30 patients. The reduction of tumor size slows greatly when OXP dose is increased to more than 100 times. In addition, increased number of treatment cycles in the IL-12 and OXP combination therapy reduced tumor size and delayed the time of tumor reaching its carrying capacity only for the 90th percentile lower responses for the 30 patients.

**Figure 6.**
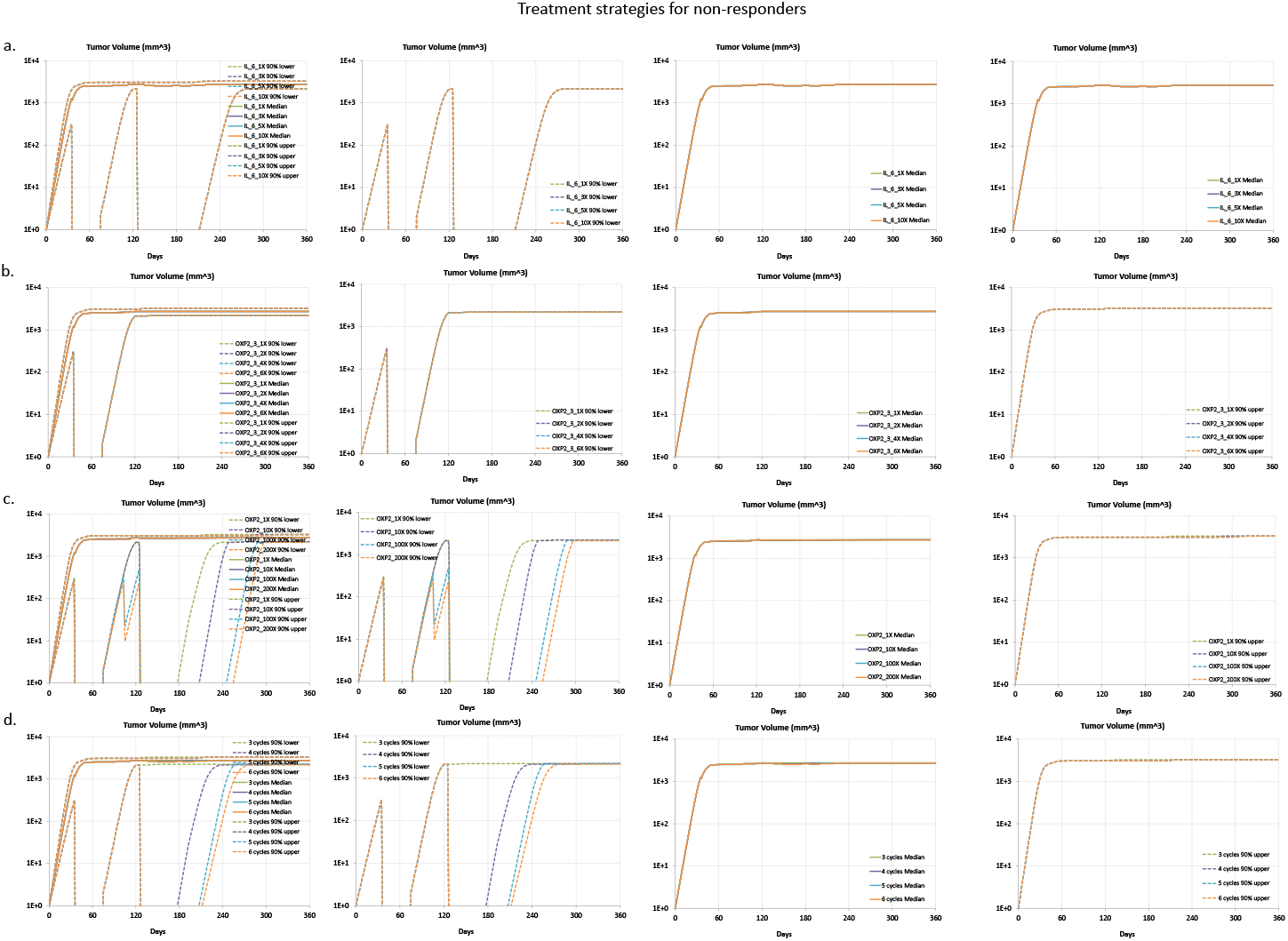
Treatment strategies for non-responders. The median, 90th percentile lower, and 90th percentile upper responses of 30 patients were sketched for each treatment strategy using 30 sets of good fits of calibrated parameters for non-responders. A sample set of parameter values used in the plots is listed in Table 2. a. Effects of increased interleukin-12 (IL-12) dose from 1 time (1X, control) to 3, 5, 10 times (3X, 5X, 10X, respectively). b. Effects of moderately increased OXP dose from 1 time (1X, control) to 2, 4, 6 times (2X, 4X, 6X, respectively). c. Effects of aggressively increased OXP dose from 1 time (1X, control) to 10, 100, 200 times (10X, 100X, 200X, respectively). d. Effects of increased number of treatment cycles from 3 cycles to 4, 5, and 6 cycles.

## Discussion and Conclusion

Developing mathematical models that represent known features of the biological system and that are calibrated to experimental studies can help improve understanding of the underlying biology targeted by drugs and enables exploring therapeutic scenarios that may be difficult or costly to test experimentally. In this paper, we developed a three-compartment mechanistic mathematical model to describe the clonal expansion of CD8^+^ T cells in a mouse model of metastatic colorectal cancer in response to a combined therapy of IL-12 plus the chemotherapy drug Oxaliplatin. Based on the collective knowledge of the underlying biology, the model represents the primary CD8^+^ T cell response under a boosting effect of IL-12 and OXP and the subsequent impact on the growth of a tumor based on the syngeneic MC38Luc1 mouse model for metastatic colorectal cancer, where the observed response was characterized by three phenotypes: responders, partial responders, and non-responders. Model parameters were calibrated against published experimental data that describes the primary response for these three phenotypes. The sensitivity analysis of parameters helped explain the differences in calibrated values of parameters between non-responders, partial-responders, and responders. To reduce the dependence of our model predictions on any single calibrated set of parameter values, we generated an ensemble of 30 parameter sets for each phenotype that provided a similar good fit against the experimental data and show the distribution in phenotypic responses for those virtual cohorts. Using the corresponding ensemble of model predictions for non-responders, numerical simulation of multiple OXP and IL-12 combination therapy suggest that aggressively increasing the dose (between 10 and 100 times of the control) of OXP will improve tumor control results while increasing the number of treatment cycles of the combined therapy can decrease the tumor size as well. We also found that only increasing the OXP dose in the combination therapy can dramatically decrease the tumor size for partial responders. Overall, these results illustrate how mechanistic models can be used to predict tumor growth response to antigen-specific immuno-chemotherapies and screen in silico for optimal therapeutic dosage and timing in treating patients with metastatic colorectal cancer.

## Ethics

The authors declare that no experiments have been performed as part of the research for this manuscript.

## Author’s contributions

DJK and QW conceived the study. DJK and QW developed the model and drafted the manuscript, ZW wrote the computer simulation code and designed the online simulators, YW performed stability analysis, QW did the numerical simulations and analyzed simulation data. All authors approved the final version of the manuscript.

## Acknowledgements

This work was supported by grants from the National Science Foundation (CAREER 1053490 to DJK), the National Cancer Institute (R01CA193473 to DJK), and the National Institute of General Medical Sciences of the National Institutes of Health grant as part of the West Virginia IDeA Network of Biomedical Research Excellence (P20GM103434 to QW). The content is solely the responsibility of the authors and does not necessarily represent the official views of the NSF, the NCI, or the National Institutes of Health. The authors would like to thank Peter Hopkins for the calibration of parameter values of functions describing relationships between IL-12 concentrations and Mif or time in days to experimental data. The authors would also like to thank Brian Crutchley, Brendan Jarrell and Vasile Stadnitchii for sensitivity analysis.

